# Six3 and Six6 jointly regulate the identities and developmental trajectories of multipotent retinal progenitor cells in the mouse retina

**DOI:** 10.1101/2023.05.03.539288

**Authors:** Alexander Ferrena, Xusheng Zhang, Rupendra Shrestha, Deyou Zheng, Wei Liu

## Abstract

Formation, maintenance, and differentiation of tissue-specific progenitor cells are fundamental tasks during organogenesis. Retinal development is an excellent model for dissecting these processes; mechanisms of retinal differentiation can be harnessed for retinal regeneration toward curing blindness. Using single-cell RNA sequencing of embryonic mouse eye cups in which transcription factor Six3 was conditionally inactivated in peripheral retinas on top of germline deletion of its close paralog Six6 (“DKO”), we identified cell clusters and then inferred developmental trajectories in the integrated dataset. In control retinas, naïve retinal progenitor cells had two major trajectories leading to ciliary margin cells and retinal neurons, respectively. The ciliary margin trajectory was directly from naïve retinal progenitor cells at G1 phase, and the retinal neuron trajectory was through a neurogenic state marked by *Atoh7* expression. Upon *Six3* and *Six6* dual deficiency, both naïve and neurogenic retinal progenitor cells were defective. Ciliary margin differentiation was enhanced, and multi-lineage retinal differentiation was disrupted. An ectopic neuronal trajectory lacking the Atoh7+ state led to ectopic neurons. Differential expression analysis not only confirmed previous phenotype studies but also identified novel candidate genes regulated by *Six3/Six6*. Six3 and Six6 were jointly required for balancing the opposing gradients of the Fgf and Wnt signaling in the central-peripheral patterning of the eye cups. Taken together, we identify transcriptomes and developmental trajectories jointly regulated by Six3 and Six6, providing deeper insight into molecular mechanisms underlying early retinal differentiation.

## Introduction

Formation, maintenance, and differentiation of tissue-specific progenitor cells are fundamental tasks during organogenesis. Retinal development in vertebrates is an excellent model for dissecting these processes, for which mechanisms can be harnessed for retinal regeneration toward curing blindness. Specification of retinal progenitor cells starts from precursor cells in the eye field at the anterior neural plate ^1^. Morphogenesis of the eye field leads to the evagination of optic vesicles, which later invaginate to form double-layered eye cups. This morphogenesis is coupled with cell-fate specification. Upon the eye-cup formation, at embryonic day 10.5 (E10.5) in mouse embryos, naïve progenitor cells for the neuroretinal epithelium and pigment epithelium are specified and compartmented to the inner and outer layer, respectively. After that, the neuroretina is further patterned along the central-peripheral axis. In central regions close to the optic stalk, naïve retinal progenitor cells start neuronal differentiation through initiating the expression of basic helix loop helix (bHLH) transcription factor gene *Atoh7* as early as E11.5 in mice ^2^, leading to the generation of retinal ganglion cells, which are born the first in the retina. Retinal ganglion cell differentiation follows a central-to-peripheral wave under the regulation of FGF signaling in chicks and mice ^3, 4^. At the far periphery of the mouse retina, naïve neural retinal progenitor cells start to express ciliary margin markers *Otx1* and *Cdon* as early as E12.5 under the regulation of Wnt/β-catenin signaling ^5, 6^. Opposing gradients of Fgf and Wnt signaling regulate the central-peripheral patterning of the eye cup ^7–9^, but it is unclear how the signaling gradients are regulated.

Retinal progenitor cells possess remarkable multipotency since they generate all types of retinal neurons and Muller glia in the retina in a conserved temporal order ^10^. Maintenance of a retinal progenitor cell pool and initiation of neuronal differentiation must be coordinated so that early- and late-born neurons are generated proportionally. Additionally, the competence of retinal progenitor cells changes over the time course ^11^. A few homeodomain and Sox family transcription factors are essential for the multipotency since their deletions in mice, two paralogs in some cases, affect the competence of retinal progenitor cells. Pax6 is required for retinal progenitor multipotency through direct transcriptional activation of bHLH transcription factors ^12^. Pax6 prevents premature activation of a photoreceptor-differentiation pathway at the periphery but regulates the multipotency of retinal progenitors in more central regions ^13^. Sox2 is required for the retinal progenitor competence and ciliary margin-neuroretina boundary in a dose-dependent manner ^14–16^. Meis1 and Meis2 are jointly required for retinal progenitor competence^17^. Despite these advances, the link between the formation and maintenance of multipotent retinal progenitor cells is largely unclear.

Our previous studies demonstrate that Six3 is required for neuroretinal specification since naïve retinal progenitor cells are absent and eye cups do not form when *Six3* is deleted at the anterior neural plate ^18, 19^. After naïve retinal progenitor cells are specified, Six3 and Six6 are jointly required for the multipotency at mid periphery and the ciliary-margin suppression at far periphery. The essential roles of Six3 and Six6 joint functions in the maintenance of multipotent retinal progenitor cells are consistent with the indispensable roles of Six3 in the specification of naïve retinal progenitor cells, providing a link between the specification and maintenance of retinal progenitor cells. Despite these advances, single-cell transcriptomes and cell trajectories regulated by Six3 and Six6 joint functions are unclear; how naïve retinal progenitor cells are regulated to differentiate into ciliary and neuronal tissues along the central-peripheral axis in the retina is still an unsolved fundamental question in the field.

Single-cell RNA sequencing (scRNA-seq) is transformative in profiling cell states and trajectories in a heterogeneous cell population and therefore is instrumental in dissecting the mechanisms underlying organogenesis. scRNA-seq has been employed to study developing tissues, including the retina ^20–25^. In scRNA-seq, individual cells are barcoded for transcriptome profiling. Tens of thousands of cells are clustered using the k-Nearest Neighbor algorithm based on the distance between individual transcriptomes. Then, differentially expressed genes (DEGs) for cell clusters are identified, elucidating expression signatures of cell states ^26^. Additionally, cell trajectories are inferred using computational approaches, including the analysis of RNA velocities ^27, 28^.

In this study, we dissect the molecular mechanisms underlying the regulation of multipotent retinal progenitor cells through transcriptional profiling of Six3 and Six6 double knockout (*Six3^F/F^;Six6^−/−^;α-Cre*) and control eye cups using scRNA-seq. We found identity signatures and developmental trajectories of naïve retinal progenitor cells in control retinas and discovered drastic defects in cell identities and trajectories upon Six3 and Six6 dual inactivation, providing deeper insight into the molecular mechanisms underlying early retinal differentiation.

## Results

### Single-cell transcriptome profiling of Six3 and Six6 double knockout and littermate control eye cups for cell clustering

To elucidate transcriptomes and cell trajectories jointly regulated by Six3 and Six6 in the maintenance and differentiation of multipotent retinal progenitor cells, we performed single-cell RNA sequencing (scRNA-seq) of Six3 and Six6 double knockout (*Six3^F/F^;Six6^−/−^;α-Cre*, referred to as DKO hereafter) and littermate control (*Six3^+/−^;Six6^+/−^*) eye cups at embryonic day 13.5 (E13.5). A reporter for the Cre activity is found in peripheral retinas starting at E10.5. In DKO retinas, Six3 protein is deleted at E11.5, and alterations of gene expression start at E12.5 and become robust at E13.5 ^7^. Therefore, the stage of E13.5 is well suited for profiling the transcriptomes regulated by Six3 and Six6. Eye cups were used in this study since the removal of lens inevitably damages ciliary margins, which are the interest of this study. Around 10,000 cells from DKO and control eye cups were captured using the Chromium fluid device for library preparation. Mapping of sequencing reads to mm10 using CellRanger revealed that control and DKO eye cups generated 10,315 cells at a depth of 21,726 reads / 2,619 genes per cell and 12,672 cells at a depth of 17,741 reads / 2,330 genes per cell, respectively.

Quality control and data analysis were further performed using the Seurat package ^26^. Cells were filtered (nFeature_RNA > 200 & nFeature_RNA < 6000 & percent.mt < 20), resulting in 9,822 and 12,146 cells for control and DKO samples, respectively. The two datasets were normalized, their top variable features were found, and the Seurat anchor-based integration method was used to combine the datasets. Cell clustering of the integrated data identified 20 clusters (Fig. 1; supp. Tab. S1). Both DKO and control cells were found in all clusters except for cluster 10 (Fig. 1A, B), which was exclusively composed of DKO cells. Notably, cluster 10 together with subsets of DKO cells in clusters 0, 3, 6 formed a DKO-specific cell population (arrowheads in Fig. 1B), indicating ectopic cell identities caused by Six3 and Six6 dual inactivation. The ectopic cell population was in G1 phase (Fig. 1B, D), as indicated by cell cycle scores determined using established cell cycle genes ^29^. Identification of the DKO-specific cell population in the scRNA-seq dataset supports drastic changes in gene expression caused by Six3 and Six6 dual inactivation described in our previous study ^7^. DKO retinas had more cells at G1 phase and fewer cells at S phase compared to control retinas (Supp. Tab. S2), indicating defects in cell cycle progression.

**Fig. 1.**
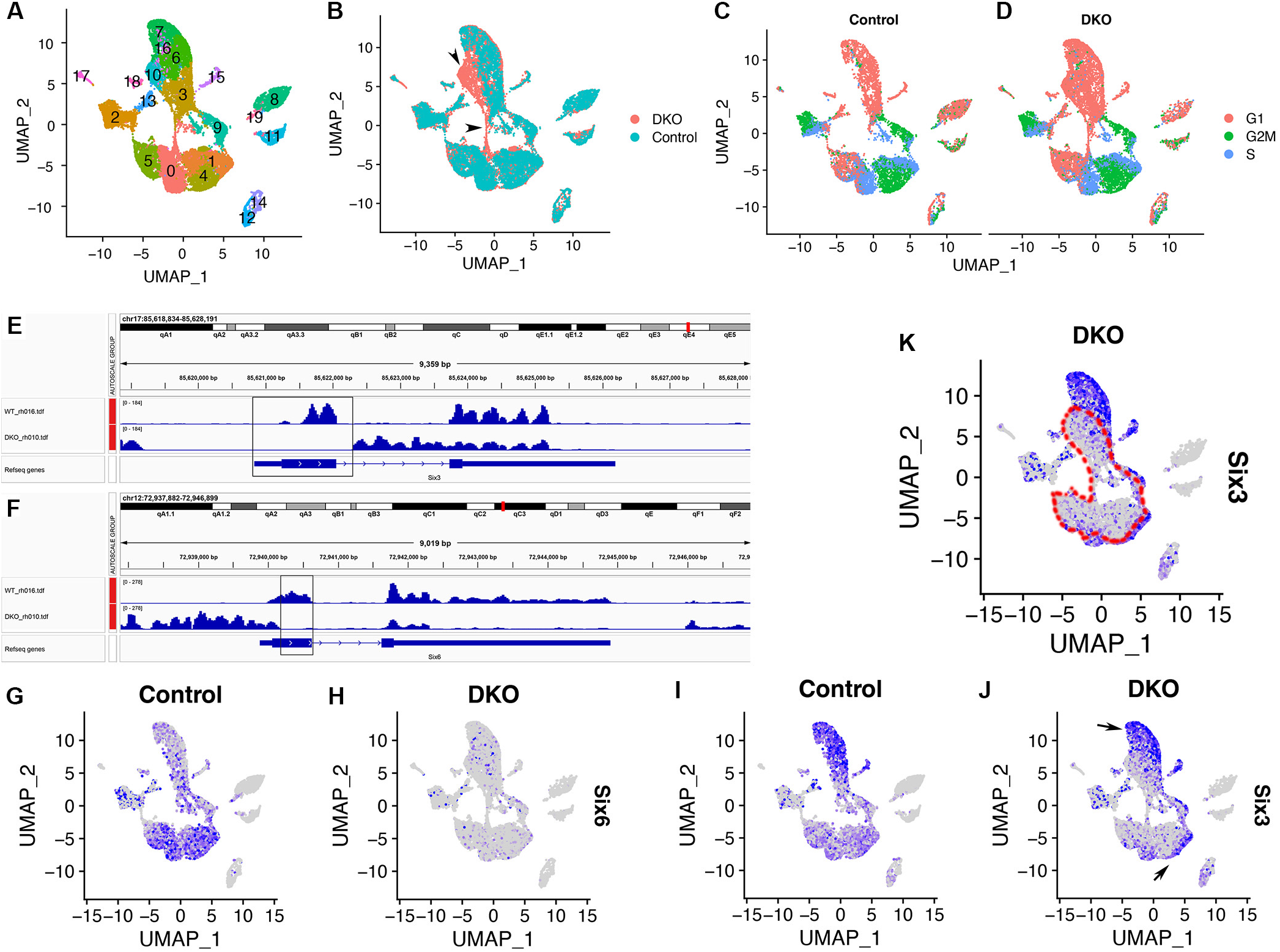
Cell clustering of the integrated scRNA-seq dataset and delineation of Six3 and Six6 dual-deficient cells in the UMAP graph. DKO and control eye cups at E13.5 were used for scRNA-seq; cell clustering was performed using the Seurat package. **(A, B)** Clustering identifies 20 cell clusters in the integrated dataset (A); cluster 10 only comprises DKO cells; a DKO-specific cell population comprising cluster 10 and subsets of clusters 0, 3, 6 is identified (arrowheads in B). **(C, D)** Cell cycle phases are labeled based cell cycle scores. **(E, F)** Genome browser views of bulk RNA-seq of DKO and control retinas reveal the deletions of targeted regions and the reductions of Six6 and Six3 mRNA coverage, which represents transcript levels. (G-H) *Six6* is widely expressed in retinal cells (G). Comparisons of *Six6* expression between control and DKO retinas reveal that *Six6* inactivation is reflected as a reduction of mRNA expression levels in the feature plot (H). (I-K) *Six3* is widely expressed in retinal cells and largely overlaps with *Six6* expression. Comparisons of *Six3* expression between control and DKO retinas reveal that conditional *Six3* inactivation is reflected as a reduction of mRNA expression levels in a well-defined area in the feature plot (outlined area in K), which area is the focus for the following study. Cells expressing *Six3* in DKO retinas are located at an outer rim (arrowheads in J) and well separated from Six3-deficient cells in the UMAP graph.

Non-retinal cells in the scRNA-seq dataset are identified based on differentially expressed genes (DEGs) between cells in one cluster and cells in all clusters (Supp. Fig. S1). Clusters 8, 19, and 11 highly expressed hemoglobin genes *Hba-X* and/or *Hba-a1*, indicating that these cells were red blood cells. Cluster 17 highly expressed *C1qb* and *Tyrobp*, indicating that these cells were white blood cells. Clusters 12 and 14 highly expressed *Cryab* and *Cryaa*, indicating that these cells were lens cells. These non-retinal clusters were well separated from other clusters in the Uniform Manifold Approximation and Projection for Dimension Reduction (UMAP) graph; they were not investigated any further.

Taken together, the scRNA-seq dataset provides an entry point in assessing transcriptomes and developmental trajectories jointly regulated by Six3 and Six6 during early retinal differentiation.

### Identification of Six3- and Six6-deficient cells in the integrated scRNA-seq dataset

In DKO retinas, *Six3* was conditionally deleted in peripheral regions by α-Cre whereas it remained in a central region; *Six6* was deleted in germline ^7^. Therefore, DKO retinas comprised both Six3-deficient and Six3-wildtype cells. We wondered whether cells with Six3 and Six6 dual deficiency in DKO retinas can be delineated in the scRNA-seq dataset so that transcriptomic changes can be correlated to Six3 and Six6 dual inactivation. In conditional *Six3* mutant mice, excision of the floxed DNA fragment in the *Six3* locus by the Cre recombinase leads to the deletion of exon 1, ablating its essential homeodomain ^30^. In *Six6* mutant mice, the DNA fragment that encodes the essential homeodomain is replaced by a drug-selection cassette in germline ^31^. As expected, bulk RNA-seq of *Six3* and *Six6* compound null retinas ^7^ demonstrates that the read coverage of targeted regions in *Six3* and *Six6* loci was absent. Importantly, the read coverage of transcripts, including the 3’ regions of mRNA that were measured by scRNA-seq, was significantly reduced (Fig. 1E, 1F). In feature plots of the scRNA-seq dataset, *Six6* was widely expressed in control retinas, but it was strongly reduced in DKO retinas (Fig. 1G, 1H). *Six3* was widely expressed in retinal cells and largely overlapped with *Six6* (Fig. 1G, 1I). When *Six3* expression in the control and DKO retinas was compared, an area with a strong reduction in expression was found in the inner regions of the UMAP graph (Fig. 1I-K), indicating that these cells were deficient for Six3. In contrast, *Six3* expression was relatively high at the outer rim in the UMAP graph (arrows in Fig. 1J). These findings indicate that Six3-deficienct and Six3- wildtype cells are well separated in the UMAP graph.

Taken together, the inactivation of *Six3* and *Six6* are reflected by reductions of mRNA levels in both bulk and scRNA-seq. Those DKO cells that showed significant reductions of *Six3* expression in the scRNA-seq dataset are deficient for both Six3 and Six6 (Fig. 1K) and therefore are the focus of this study. The DKO-specific cell population is a subset of Six3- and Six6-deficient cells (Fig. 1B, 1K).

### Retinal progenitor cells comprise naïve and neurogenic populations in control retinas and both cell populations are defective upon Six3 and Six6 dual deficiency

We then assessed the identities of retinal cell clusters using known gene markers and cluster-specific DEGs identified this study. In control retinas, clusters 0, 1, 4, 5 highly expressed *Ccnd1*, *Sfrp2*, and *Vsx2*, (Fig. 2A, 2C, 2E), which are well established gene markers for naïve retinal progenitor cells. These clusters also highly expressed *Crabp2* and *Dapl1* based on our scRNA-seq data (Fig. 2G, 2I). Clusters 0, 1, 4, and 5 were grossly aligned with cell cycle phases (Fig. 1A, C). In contrast to these naïve retinal progenitor cells, clusters 9 and 3 highly expressed *Atoh7* (Fig. 2K), indicating their neurogenic identities. Clusters 9 and 3 were also grouped along cell cycle phases (compare Fig. 2K with Fig. 1C). A small portion of Atoh7+ retinal progenitor cells also expressed *Ccnd1*, *Sfrp2*, and *Vsx2* (arrowheads in Fig. 2A, 2C, 2E, 2K), indicating that these cells were in transition from the naïve retinal progenitor state to the neurogenic progenitor state. *Gadd45a* expression largely overlapped with *Atoh7* expression in neurogenic retinal progenitor cells (Fig. 2K, 2M). A subset of Atoh7+ neurogenic retinal progenitor cells also expressed *Gm14226*, *Neurod1*, and *Bhlhe22* (Fig. 2K, 2O, 2Q, 2S).

**Fig. 2.**
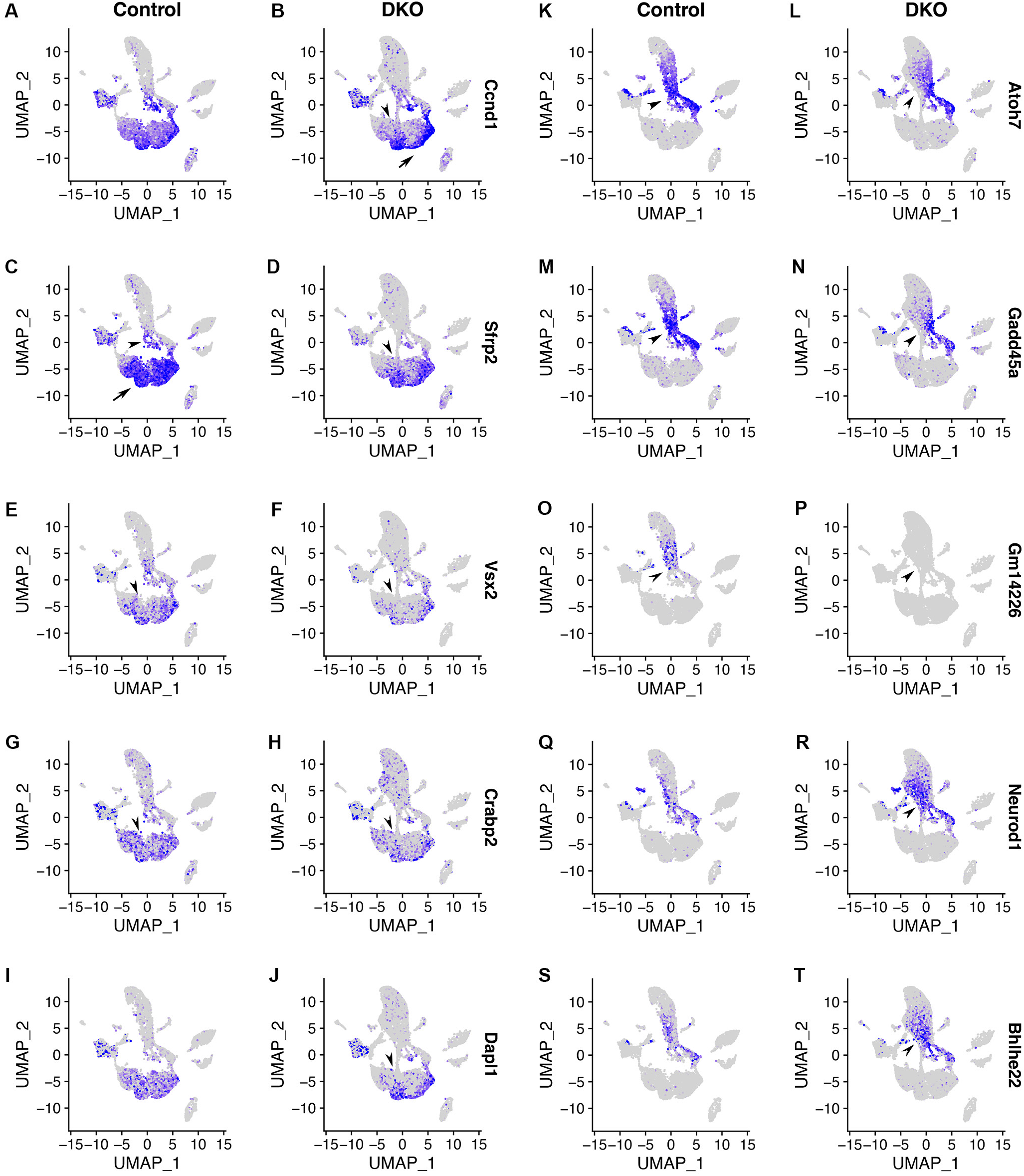
Retinal progenitor cells comprise naïve and neurogenic populations in control retinas and both cell populations are defective upon Six3 and Six6 dual deficiency. **(A-J)** In control retinas, clusters 0, 1, 4, 5 differentially express *Ccnd1*, *Sfrp2*, and *Vsx2* (A, C, E, respectively), indicating that these cells are naïve retinal progenitor cells. These naïve retinal progenitor cells also differentially express gene markers *Crabp2* and *Dapl1* that were identified in this study (G, I). Cluster 2 also express these naïve retinal progenitor cell markers, but cluster 2 is marked by negative gene markers (see also supp. Fig. S1) and have lower values for nCount_RNA and nFeature_RNA (see also supp. Fig. S3), indicating the differences between cluster 2 and clusters 0, 1, 4, 5. In Six3- and Six6-deficient cells, the expression of these naïve retinal progenitor cell markers is significantly reduced (B, D, F, H, J, respectively; please use the outlined area in Fig. 1K to delineate Six3 and Six6 dual deficiency). **(K-T)** In the control retinas, clusters 9 and 3 differentially express *Atoh7* (K), indicating that these cells are neurogenic retinal progenitor cells. Naïve and neurogenic retinal progenitor cells are separated in the UMAP graph and marked by distinct gene markers (A, K), confirming that they are two cell populations. Neurogenic retinal progenitor cells also express gene markers *Gadd45a* and *Gm14226* identified in this study (M, O). Subsets of neurogenic retinal progenitor cells also express *Neurod1* and *Blhlhe22* (Q, S). In Six3- and Six6-deficient cells, the expression of *Atoh7*, *Gadd45a*, and *Gm14226* is significantly downregulated (L, N, P, respectively), whereas the expression of *Neurod1* and *Blhlhe22* is significantly upregulated (R, T).

Upon Six3 and Six6 dual deficiency, the expression of *Ccnd1*, *Sfrp2*, *Vsx2*, *Crabp2*, *Dapl1*, *Atoh7*, *Gadd45a*, and *Gm14226* was strongly reduced or nearly absent (arrowheads in Fig. 2B, 2D, 2F, 2H, 2J, 2L, 2N, 2P; Six3 and Six6 dual deficiency is outlined in Fig. 1K), whereas the expression of *Neurod1* and *Bhlhe22* was significantly increased (arrowheads in Fig. 2R, 2T). Neurod1 is a pan-amacrine transcription factor in mouse retina ^32^. Bhlhe22 (also known as Bhlhb5) overlaps with Neurod1 and is required for the subtype development of retinal amacrine and bipolar cells in mice ^32^. The upregulation of *Neurod1* and *Bhlhe22* in the scRNA-seq dataset is consistent with increased number of amacrine cells in DKO retinas described in our previous study ^7^. Taken together, naïve and neurogenic retinal progenitor cells form two cell populations in the UMAP graph; both cell populations are defective upon Six3 and Six6 dual deficiency.

We noted that cluster 2 also expressed naïve retinal progenitor cell markers (Fig. 2A-2J) and cluster 13 also expressed neurogenic markers (Fig. 2K-2N) for in both control and DKO retinal cells, but these two clusters were separated from main groups of retinal progenitor cells. Compared to clusters 5, 0, 4, 1, 9, and 3, clusters 2 and 13 were defined by negative gene markers, including *Kcnq1ot1*, *mt-Co1*, *mt-Nd2*, and *mt-Nd4* (Supp. Fig. S2). Interestingly, clusters 2 and 13 had lower values for nCount_RNA, nFeature_RNA, and percent.mt (Supp. Fig. S3). Therefore, clusters 2 and 13 were separate retinal cell populations that had lower counts in RNA molecules and genes, perhaps due to lower rates of RNA capture by unknown technical reasons. Alternatively, the low counts in RNA molecules and genes in clusters 2 and 13 may reflect uncharacterized retinal cells that have a lower number of expressed genes. In support of this notion, nFeature_RNA varied significantly along cell clusters; clusters 8 and 11, which were red blood cells, also had low counts in nFeature_RNA. Further studies are needed to determine why clusters 2 and 13 are separated from main groups of retinal cells in the UMAP graph.

### Early retinal neurons follow the Atoh7+ neurogenic state in the UMAP graph of control retinas and retinal differentiation is defective upon Six3 and Six6 dual deficiency

We next assessed the differentiation of retinal ganglion cells, amacrine cells, and photoreceptor cells in the scRNA-seq dataset. In control cells, clusters 6 and 3 were marked by the expression of *Pou4f2*, *Dlx1*, *Igfbp5*, *Dlx2*, and *Isl1* (Fig. 3A, 3C, 3E, 3G, 3I; cluster information is in Fig. 1A), which are known to regulate retinal ganglion cell differentiation. The expression of these gene markers overlapped with *Atoh7* expression in cluster 3 (compare Fig. 3A, 3C, 3E, 3G, 3I with Fig. 2K). Additionally, *Isl1* expression marked clusters 6, 3 and all the way to clusters 7 and 16 (Fig. 3I). Cells at the end of this branch highly expressed *Sncg* (Fig. 3K), indicating that Sncg+ cells were retinal ganglion cells at advanced differentiation stages. The expression patterns of *Ccnd1*, *Atoh7*, *Pou4f2*, *Isl1*, and *Sncg* in the UMAP graph revealed progressively-advancing states during retinal ganglion cell differentiation in control retinas (Fig. 2A, 2K, Fig. 3A, 3I, 3K). Therefore, the UMAP captures a progression of cell states starting from naïve retinal progenitors, through neurogenic progenitors, and to retinal ganglion cells at advanced stages.

**Fig. 3.**
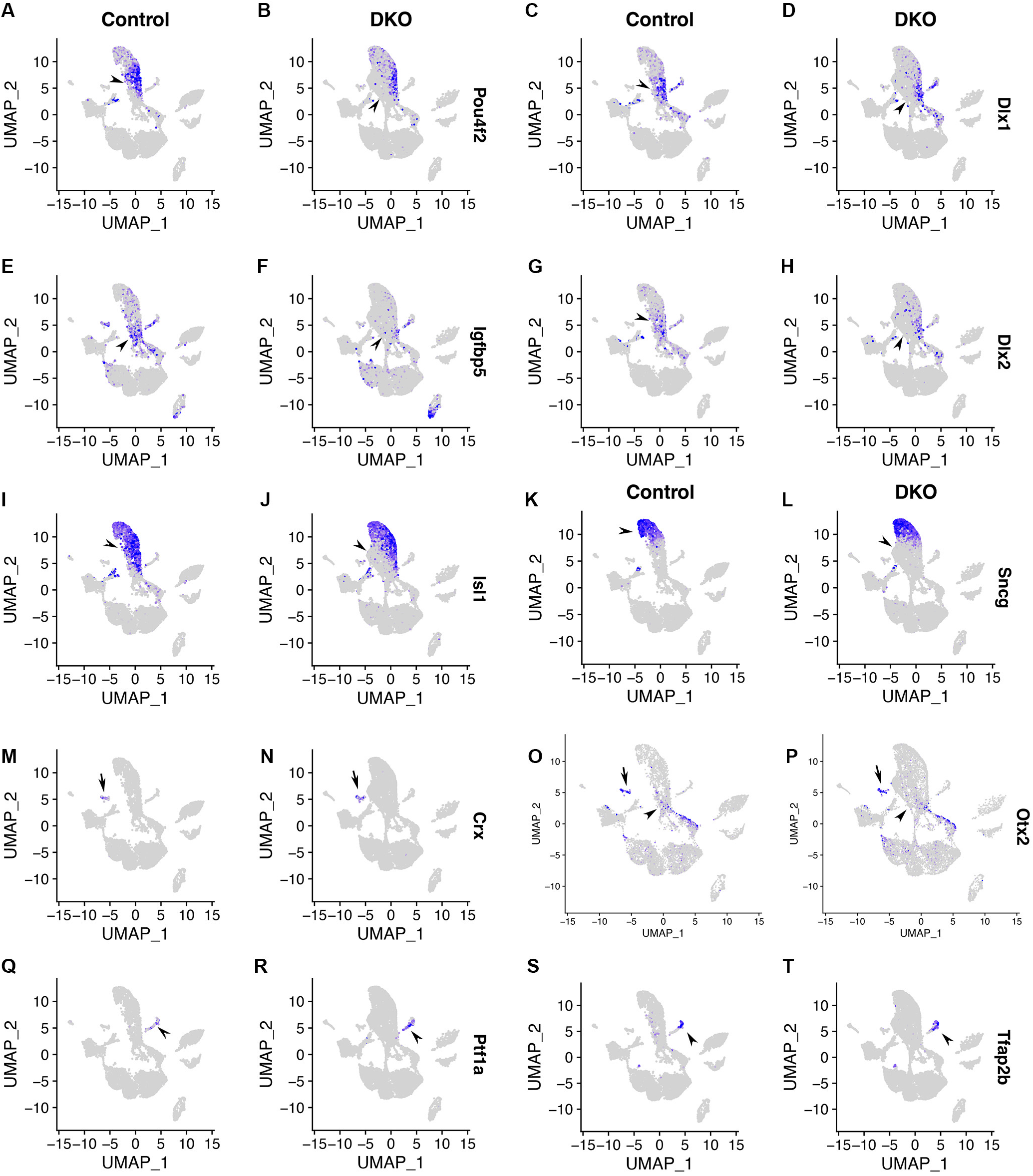
Early retinal neurons follow the Atoh7+ neurogenic state in the UMAP graph of control retinas and retinal differentiation is defective upon Six3 and Six6 dual deficiency. **(A-L)** In control retinas, retinal ganglion cell markers *Pou4f2*, *Dlx1*, *Igfbp5*, *Dlx2*, and *Isl1* are expressed in cells adjacent to or partially overlapped with Atoh7+ cells in the UMAP graph (A, C, E, G, I, respectively). Retinal ganglion cell marker *Sncg* is expressed in cells at the end of this branch (K). Expression patterns of *Atoh7*, *Pou4f2*, *Isl1*, and *Sncg* in the UMAP graph indicate a progression of gene expression starting from neurogenic retinal progenitor cells toward retinal ganglion cells at advanced stages. In Six3- and Six6-deficient cells, the expression of *Pou4f2*, *Dlx1*, *Igfbp5*, *Dlx2*, and *Isl1* is significantly downregulated (B, D, F, H, J, respectively). Cells expressing Sncg in DKO retinas are in Cre-negative areas since Six3 remains (please compare Fig. 3K, 3L with Fig. 1K). **(M-P)** Early photoreceptor cells are marked by the expression of *Crx* and *Otx2* (arrows in M-P). In control retinas, *Otx2* is also expressed in neurogenic retinal progenitor cells, but its expression is downregulated in neurogenic retinal progenitor cells upon Six3 and Six6 dual deficiency (arrowheads in O, P). **(Q-T)** Amacrine and horizontal cells are marked by *Ptf1a* and *Tfap2b* expression, and the expression of *Ptf1a* and *Tfap2b* is up- regulated in DKO retinas.

In Six3- and Six6-deficient retinal cells, however, the expression of *Pou4f2*, *Dlx1*, *Igfbp5*, *Dlx2*, and *Isl1* was absent or drastically reduced (Fig. 3B, 3D, 3F, 3H, 3J). Remaining Sncg+ cells in DKO retinas were in regions where *Six3* expression remained (compare Fig. 3K, 3L with Fig. 1I- K). These findings indicate that retinal ganglion cell differentiation is defective upon Six3 and Six6 dual deficiency.

The differentiation of photoreceptor and amacrine cells was also defective upon Six3 and Six6 dual deficiency. In control retinal cells, cluster 18 was marked by the expression of *Crx* and *Otx2* (Fig. 3M, 3O). Interestingly, *Otx2* was also expressed in a subset of neurogenic retinal progenitor cells in controls (arrowhead in Fig. 3O), and *Otx2* expression in neurogenic retinal progenitor cells was reduced upon Six3 and Six6 dual deficiency (arrowhead in Fig. 3P), indicating defects in photoreceptor cell differentiation caused by Six3 and Six6 dual inactivation. Cluster 15 was marked by the expression of *Ptf1a* and *Tfap2b* in control cells (Fig. 3Q, 3S), indicating that these cells were horizontal and/or amacrine cells. The expression of *Ptf1a* and/or *Tfap2b* was increased in DKO cells (Fig. 3R, 3T), indicating enhanced differentiation toward horizontal/amacrine cells. Collectively, early retinal differentiation is defective upon Six3 and Six6 dual deficiency.

### The DKO-specific cell population ectopically expresses gene markers for the forebrain and amacrine cells

The DKO-specific cell population included cluster 10 and subsets of cells in clusters 0, 3, and 6 (arrowheads in Fig. 1B); they barely express neurogenic progenitor marker *Atoh7* (Fig. 2K) and retinal ganglion cell markers *Pou4f2*, *Dlx1*, *Igfbp5*, *Dlx2*, and *Isl1* (Fig. 3B, 3D, 3F, 3H, 3J). To determine the identities of this DKO-specific cell population, we identified DEGs between cluster 10 (DKO-specific) and control cluster 6. Cluster 10 and control cluster 6 were compared since there were no control cells in cluster 10 and control cluster 6 was close to cluster 10 in the UMAP. Furthermore, both PCA of cluster averages and hierarchical clustering of correlation matrix showed that cluster 10 is closest with cluster 6. We also identified DEGs between DKO and control cells in clusters 3 and 9, and DEGs between DKO and control cells in clusters 0, 1, 4, 5 (see also Fig. 8). Using feature plotting of the top DEGs, we identified genes that exhibited specific expression in the DKO-specific cell population. *Foxg1* and *Epha3*, two genes that are strongly expressed in the forebrain and weakly expressed in the retina in E14.5 mouse embryos (based on data in the genepaint database), were significantly upregulated in the DKO-specific cell population (Fig. 4A-4D). Similarly, gene markers *Dmrta2*, *Sorbs2*, *Enc1*, *Uncx*, *St18*, *Tac1*, *Lhx9*, and *Samd5* were also significantly up-regulated (Fig. 4E-T). In E14.5 mouse embryos, *Dmrta2*, *Sorbs2*, *Enc1*, *Tac1*, *Lhx9*, and *Samd5* are highly expressed in the forebrain; *Dmrta2* is modestly expressed in the retina; *Sorbs2*, *Enc1*, *Tac1*, and *Lhx9* are weakly expressed in the retina; *Samd5* is undetectable in the retina (based on data in the genepaint database). In our scRNA-seq dataset, the expression of *Dmrta2*, *St18*, *Tac1*, *Lhx9*, and *Samd5* were barely detectable in control retinas (Fig. 4E, 4M, 4O, 4Q, 4S). Besides these markers, *Tfap2c*, a gene marker that is expressed in postmitotic amacrine cells in mice ^33^, was significantly up-regulated in a subset of the DKO-specific cell population (Fig. 4U, 4V). The expression of *Cdkn1c* (also known as *P57Kip2*), a negative regulator of cell proliferation, was also up-regulated in a subset of the DKO-specific cell population (Fig. 4W, 4X). Additionally, the expression of *Foxg1*, *Epha3*, *Dmrta2*, *Enc1*, *Tac1*, *Samd5*, and *Tfap2c* were up-regulated in both the DKO-specific cell population and naïve retinal progenitor cells. Taken together, these findings indicate that gene markers for the forebrain and amacrine cells are significantly up-regulated in the DKO-specific cell population.

**Fig. 4.**
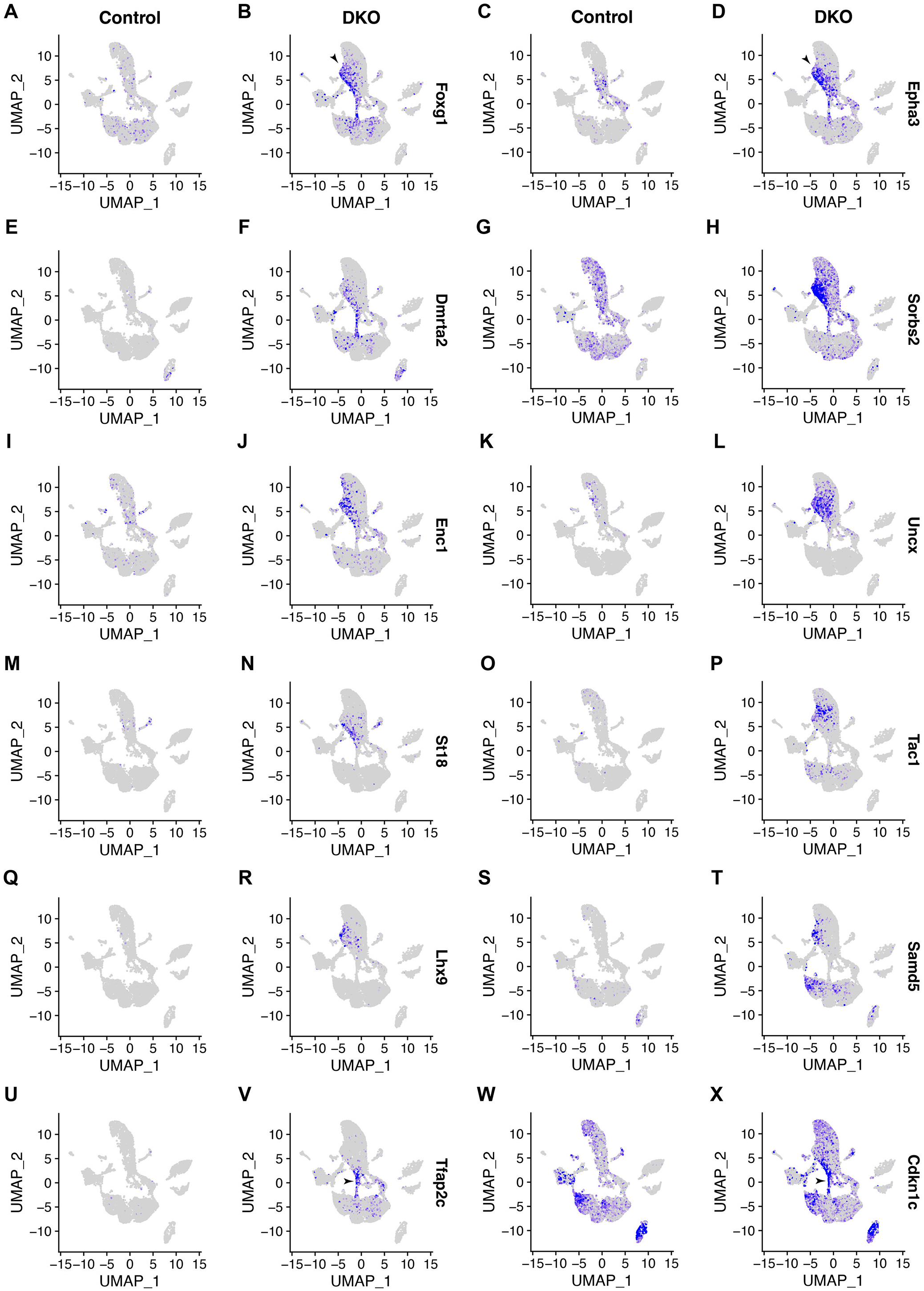
The DKO-specific cell population ectopically expresses gene markers for the forebrain and amacrine cells. **(A-T)** Feature plots of DEGs that are differentially up-regulated in the DKO-specific cell population. In control retinas, these gene markers are either weakly expressed or undetectable, but they are highly expressed in the forebrain (see also the genepaint database). In the DKO-specific cell population, these gene markers are significantly upregulated. The expression of *Foxg1*, *Epha3*, *Dmrta2*, *Tac1*, and *Samd5* is also significantly upregulated in naïve retinal progenitor cells upon Six3 and Six6 dual deficiency. **(U, V)** Amacrine cell marker *Tfap2c* is upregulated in some DKO-specific cells and naïve retinal progenitor cells upon Six3 and Six6 dual deficiency. **(W, X)** Cell cycle inhibitor *Cdkn1c* is upregulated in a subset of the DKO-specific cell population (arrowhead in X).

### Ciliary margin cells are at an edge of the G1-phase naïve retinal progenitor cell cluster in control retinas and this cell population is expanded upon Six3 and Six6 dual deficiency

We next assessed ciliary margin cells in the scRNA-seq dataset. In control retinas, cells at an edge of cluster 5, the G1-phase naïve retinal progenitor cell cluster, were marked by the expression of *Gja1*, *Otx1*, *Trpm1*, *Apoe*, and *Dct* (Fig. 5). *Gja1* and *Otx1* are expressed in ciliary margins of E14.5 mouse embryos (genepaint); a homolog of *Trpm1* in zebrafish is expressed in ciliary margins ^34^. Our data indicates that *Apoe* and *Dct* were expressed in a subset of ciliary margins (Fig. 5I, 5K). In contrast to these ciliary margin markers, the expression of naïve retinal progenitor cell markers *Ccnd1*, *Sfrp2*, and *Vsx2* was lower in this region (Fig. 2A, 2C, 2E). In DKO retinas, the ciliary margin cell population was expanded toward the naïve retinal progenitor cell population (Fig. 5). Taken together, the ciliary margin cell population is at an edge of the G1- phase naïve retinal progenitor cell cluster in control retinas and this population is expanded upon Six3 and Six6 dual deficiency.

**Fig. 5.**
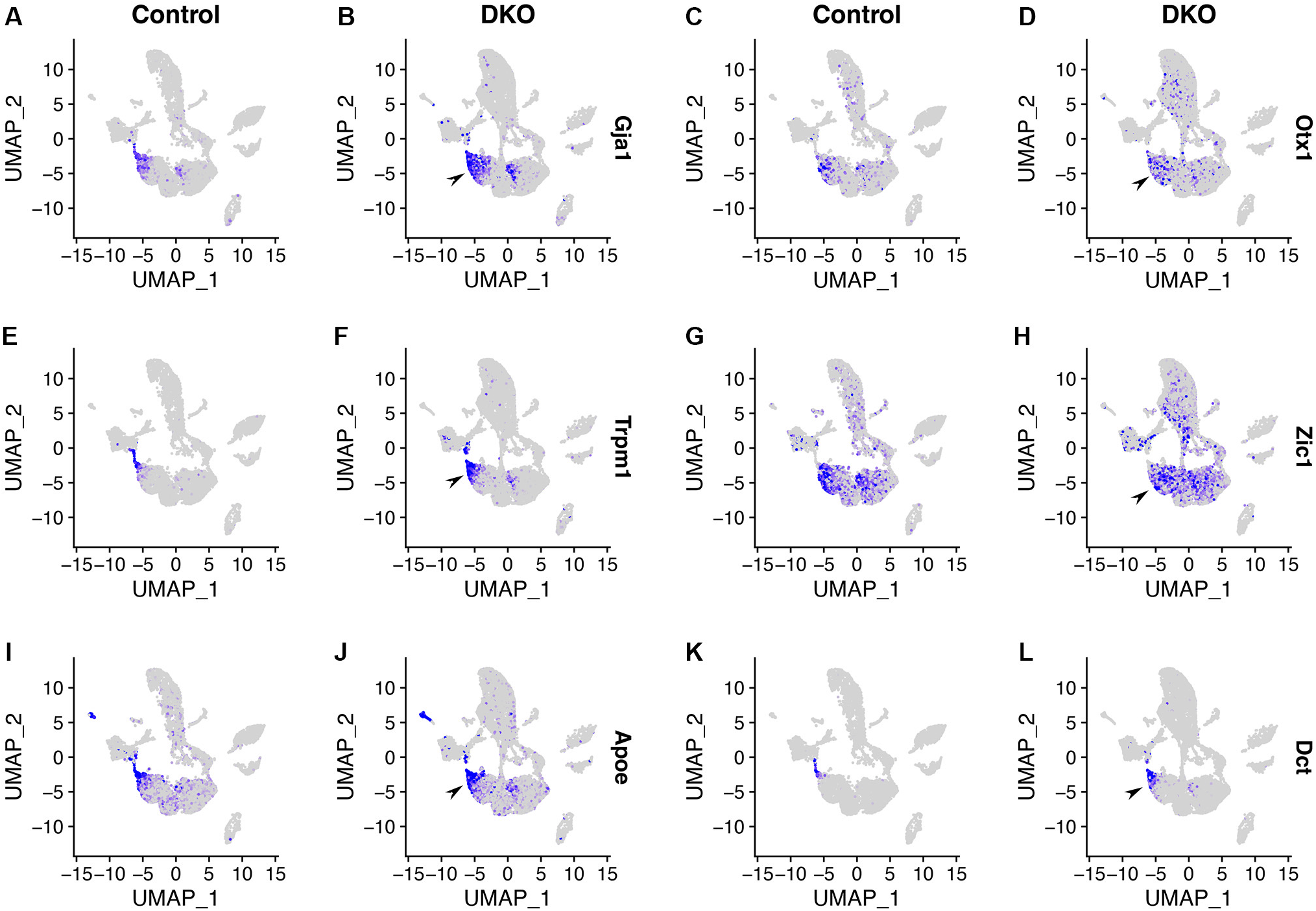
Ciliary margin cells are at an edge of the G1-phase naïve retinal progenitor cell cluster in control retinas and this cell population is expanded upon Six3 and Six6 dual deficiency. **(A-L)** In control retinas, ciliary margin cells are marked by the expression of *Gja1*, *Otx1*, *Trpm1*, *Zic1*, *Apoe*, and *Dct* in the UMAP graph (A, C, E, G, I, K, respectively). In DKO retinas, these gene markers are significantly upregulated (B, D, F, H, J, L, respectively).

### Regulators of the Wnt pathway, Fgf pathway, and actin cytoskeleton are affected by Six3 and Six6 dual deficiency

Our differential expression analysis between DKO and control cells identified defects in numerous signal transduction pathways upon Six3 and Six6 dual deficiency. In the Wnt pathway, the expression of *Fzd1* and *Axin2* was widely up-regulated in dual deficient cells (compare Fig. 6A-6D with Fig. 1I-1K). In contrast, the expression of *Wls* and *Rspo3* was significantly upregulated in clusters 5 and 10, respectively (Fig. 6F, 6H). In addition, *Wnt16* expression was up-regulated in cluster 5 whereas *Fzd5* expression was down-regulated in naïve retinal progenitor cells (Fig. 6J, 6L). In the FGF pathway, the expression of *Fgf9* and *Fgf15* was strongly downregulated in naïve retinal progenitor cells, with stronger reduction in *Fgf9* expression (Fig. 6N, 6P). The expression of *Filip1*, which promotes the degradation of filamin A and therefore remodels the actin cytoskeleton ^35^, was drastically up-regulated (compare Fig. 6R with Fig. 1I-1K). The expression of *Ezr*, which serves as an intermediate between the plasma membrane and the actin cytoskeleton ^36^, was upregulated in naïve retinal progenitor cells (Fig. 6T). These findings indicate that regulators of the Wnt pathway, Fgf pathway, and actin cytoskeleton are affected by Six3 and Six6 dual deficiency.

**Fig. 6.**
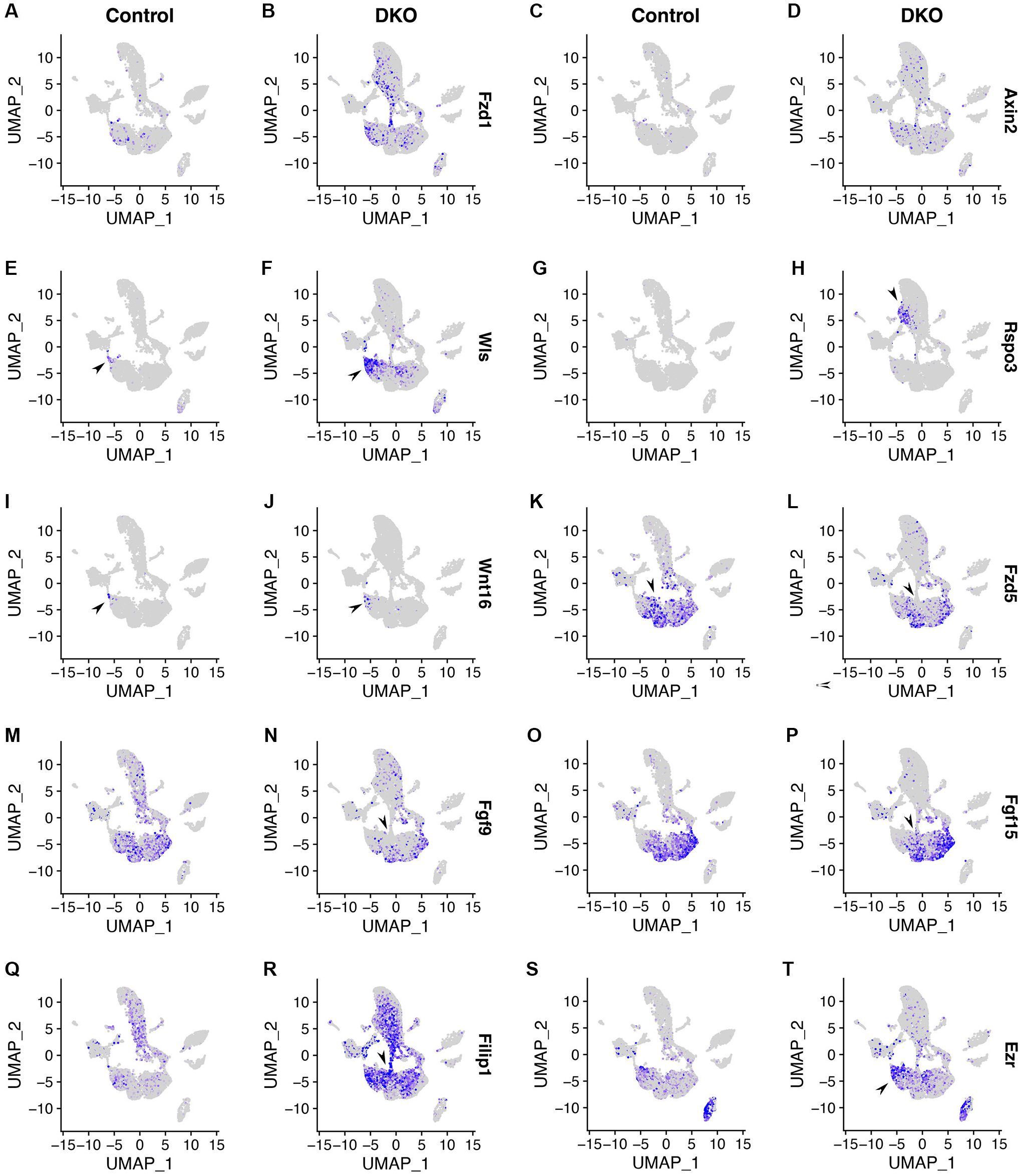
Regulators of the Wnt pathway, Fgf pathway, and actin cytoskeleton are affected by Six3 and Six6 dual deficiency. **(A-L)** In control retinas, *Fzd1*, *Wls,* and *Wnt16* are expressed in areas corresponding to ciliary margins in the UMAP graph (A, E, I, respectively); *Axin2* expression is low (C); *Rspo3* expression is undetectable (G); *Fzd5* is widely expressed in areas corresponding to naïve retinal progenitor cells. In DKO retinas, *Fzd1*, *Axin2*, *Wls*, *Rspo3*, and *Wnt16* are significantly upregulated (B, D, F, H, J) whereas *Fzd5* is significantly downregulated (L). **(M-P)** In control retinas, *Fgf9* and *Fgf15* are highly expressed in areas corresponding to naïve retinal progenitor cells in the UMAP graph. In Six3- and Six6-deficient cells, *Fgf9* and *Fgf15* are significantly downregulated (N, P; Six3- and Six6-deficient areas are outlined in Fig. 1K). **(Q-T)** Actin cytoskeleton regulators *Filip1* and *Ezr* are significantly upregulated upon Six3 and Six6 dual deficiency.

### Validation of major DEGs using in situ hybridization

To validate DEGs identified using scRNA-seq, we performed in situ hybridization of DKO and control embryos at E13.5 (Fig. 7). In control retinas, *Gja1* was strongly expressed at ciliary margins. In DKO retinas, *Gja1* expression was expanded to peripheral retinas. In control retinas, *Dct* was expressed at the retina pigment epithelium and at the tip of ciliary margins. In DKO retinas, *Dct* expression was drastically upregulated in peripheral retinas. In control retinas, *Wls* was weakly expressed at the tip of ciliary margins. In DKO retinas, *Wls* expression was significantly upregulated in peripheral retinas. In control retinas, *Rspo3* did not show any specific expression above the background level. In DKO retinas, *Rspo3* expression was significantly upregulated in peripheral retinas. Therefore, major DEGs identified using scRNA-seq are confirmed by expression analysis.

**Fig. 7.**
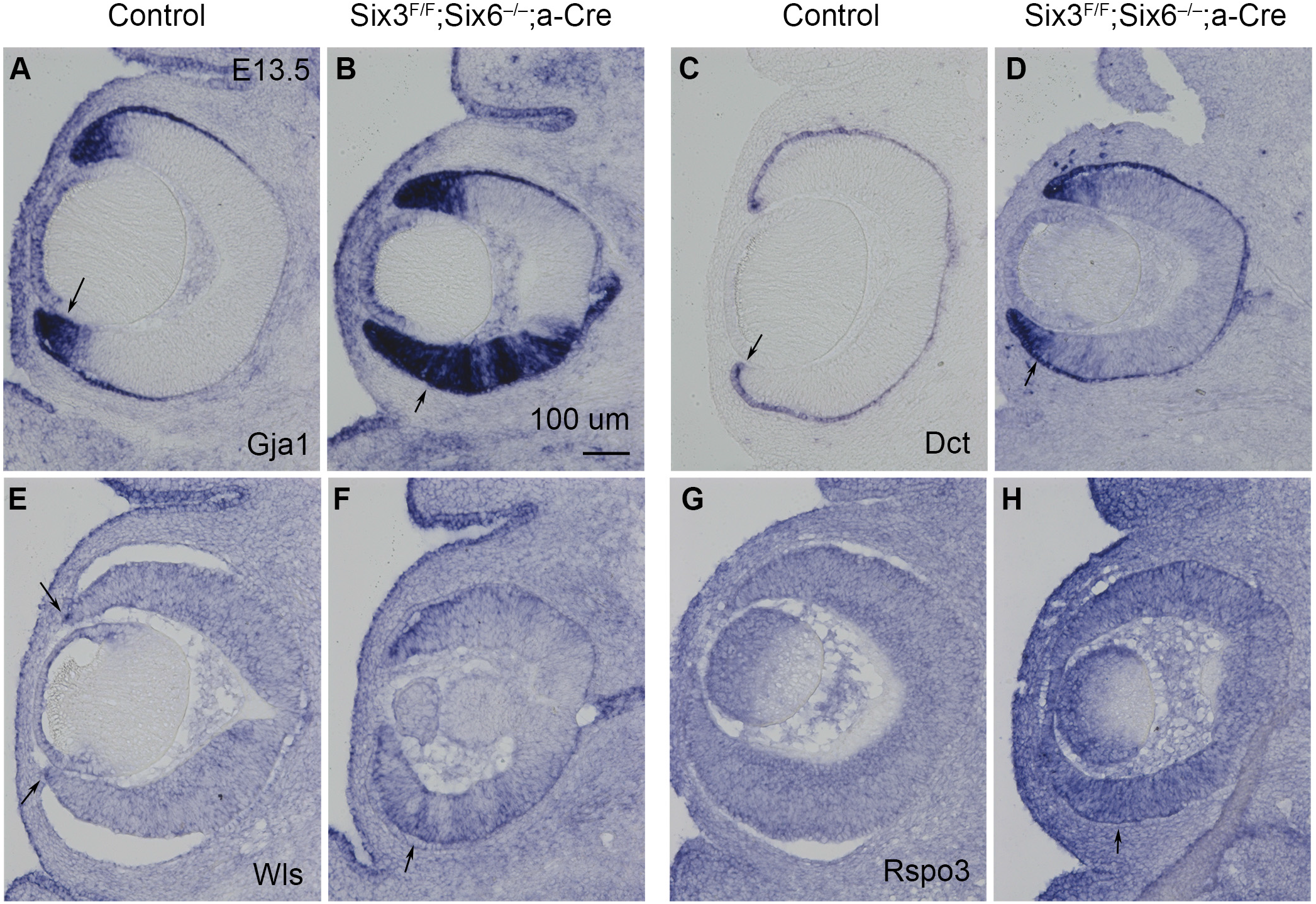
Validation of major DEGs using *in situ* hybridization. **(A, B)** *Gja1* expression is drastically upregulated in peripheral retinas upon Six3 and Six6 dual deficiency. **(C, D)** *Dct* expression is significantly upregulated in peripheral retinas upon Six3 and Six6 dual deficiency. **(E, F)** *Wls* expression is significantly upregulated in peripheral retinas upon Six3 and Six6 dual deficiency. **(G, H)** *Rspo3* expression is significantly upregulated in peripheral retinas upon Six3 and Six6 dual deficiency. Scale bar, 100 μm.

### Identification of enriched GO terms in DEGs caused by Six3 and Six6 dual inactivation

To annotate the functions of DEGs between DKO and control cells in clusters 0, 1, 4, 5, DEGs between DKO and control cells in clusters 3 and 9, and DEGs between cluster 10 (DKO-specific) and control cluster 6, we performed GO term analysis. Top enriched GO terms in DEGs (188, padj. <0.01) between DKO and control cells in clusters 0, 1, 4, 5 included eye development, axonogenesis, forebrain development, epithelial cell proliferation, cell junction assembly, and Wnt signaling pathway (Fig. 8A). Top GO terms in DEGs (344, padj. <0.01) between DKO and control cells in clusters 3 and 9 included axonogenesis, sensory system development, forebrain development, and mitotic cell cycle phase transition (Fig. 8B). Top GO terms in upregulated DEGs (287, padj. < 0.01) between cluster 10 (DKO-specific) and control cluster 6 include axonogenesis, forebrain development, synapse organization, and telencephalon development (Fig. 8C). Top GO terms in downregulated DEGs (528, padj. < 0.01) between cluster 10 (DKO-specific) and control cluster 6 included axonogenesis, regulation of cellular amide metabolic process, positive regulation of protein localization, regulation of neurogenesis, and neuron death (Fig. 8D). Taken together, DEGs in these three groups are enriched with both related and distinct GO terms.

**Fig. 8.**
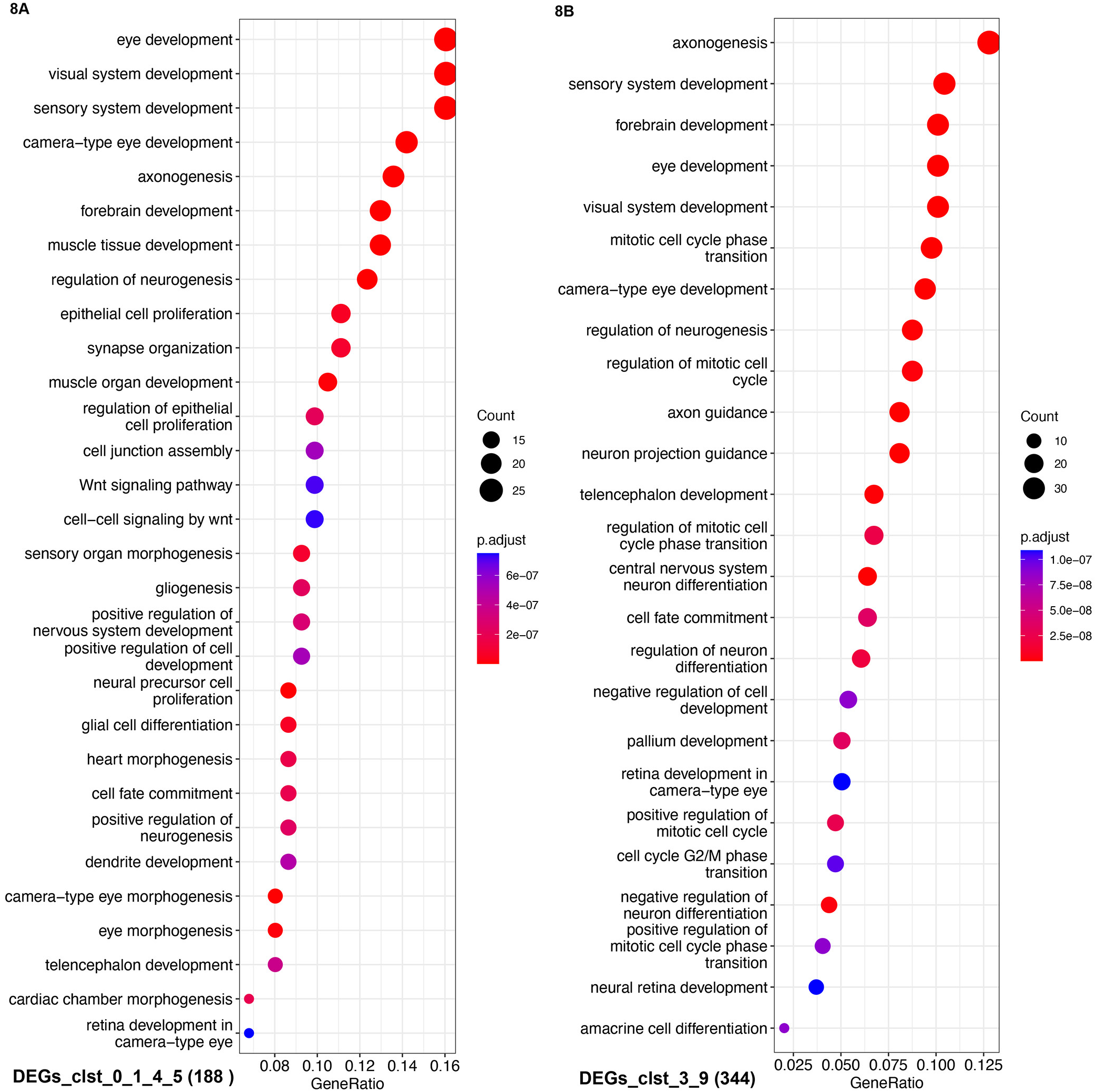
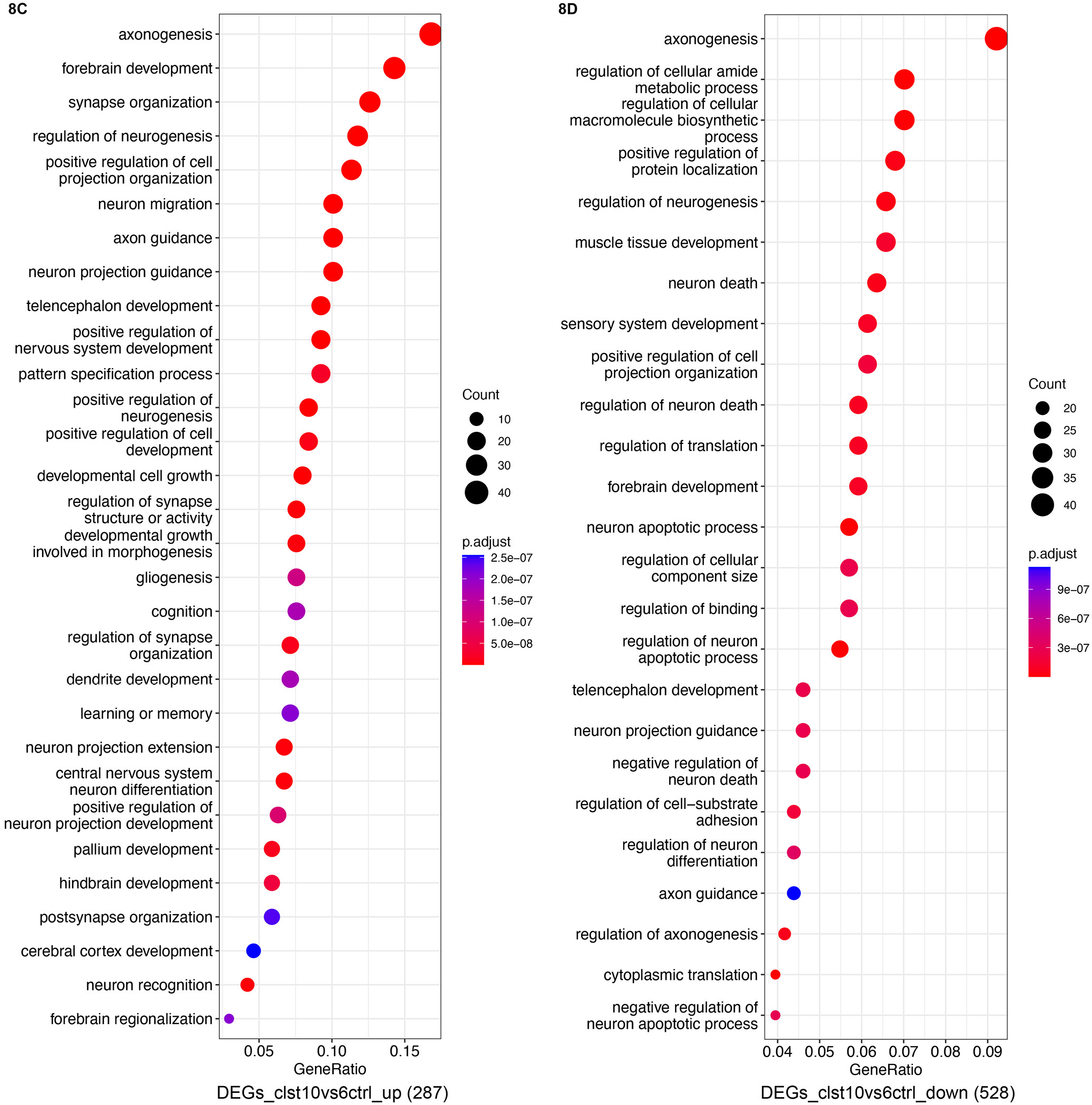
Identification of enriched GO terms in DEGs caused by Six3 and Six6 dual inactivation. **(A)** Enriched GO terms in DEGs caused Six3 and Six6 dual inactivation in clusters 0, 1, 4, and 5. **(B)** Enriched GO terms in DEGs caused Six3 and Six6 dual inactivation in clusters 3 and 9. **(C)** Enriched GO terms in upregulated DEGs between DKO-specific cluster 10 and control cluster 6. **(D)** Enriched GO terms in downregulated DEGs between DKO-specific cluster 10 and control cluster 6.

### Developmental trajectories of naïve retinal progenitor cells are altered upon Six3 and Six6 dual deficiency

To elucidate developmental trajectories of control and DKO retinal cells, we analyzed RNA velocities of retinal cells in the integrated dataset using the scVelo package ^28^. In control retinas, naïve retinal progenitor cells had two major trajectories: one trajectory was toward ciliary margin cells in cluster 5, and the other trajectory was toward neurons in cluster 7 through a Atoh7+ neurogenic state (Fig. 9A). In DKO retinas, the trajectory toward ciliary margin cells was enhanced; the trajectory toward neuronal differentiation was defective. An ectopic trajectory lacking a Atoh7+ state led to ectopic neurons in cluster 10 (Fig. 9B; see also Fig. 1B, 2L). Therefore, Six3 and Six6 jointly regulate developmental trajectories of naïve retinal progenitor cells in the mouse retina.

**Fig. 9.**
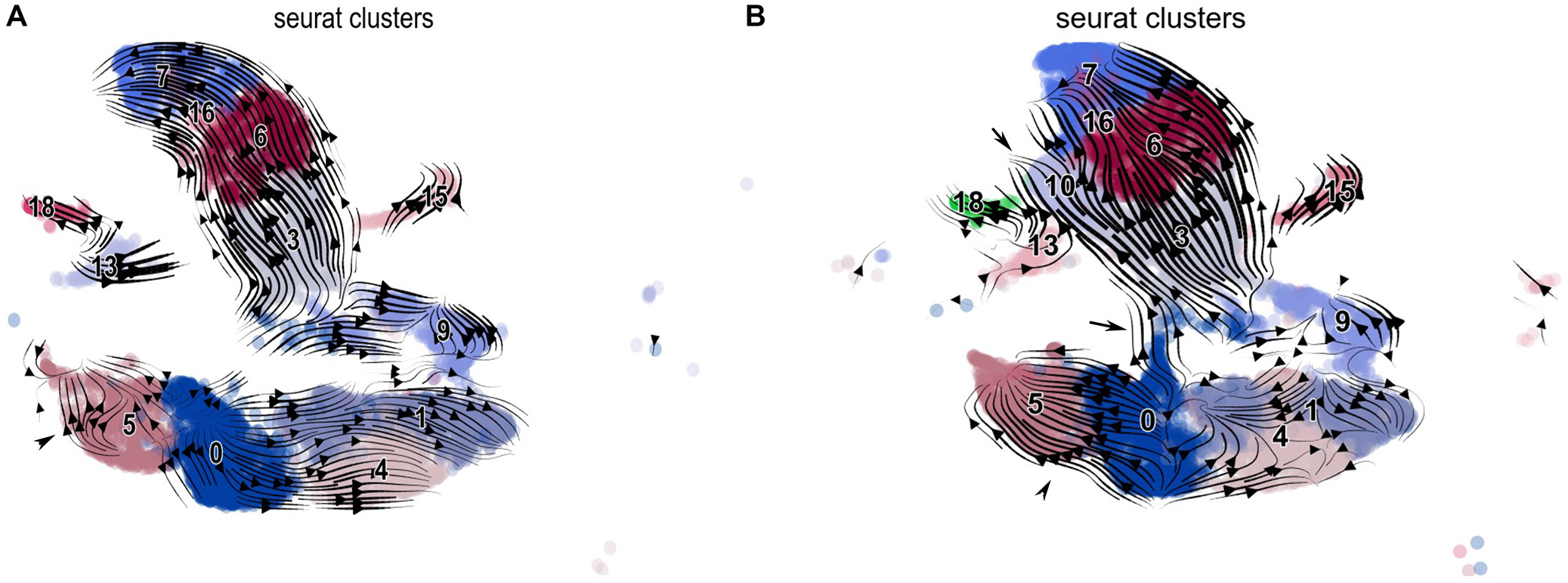
Developmental trajectories of naïve retinal progenitor cells are altered upon Six3 and Six6 dual deficiency. Non-retinal cells and clusters 2 (cluster 2 has low values for nCount_RNA and nFeature_RNA) were removed before the cell trajectory analysis. **(A, B)** In control retinas, naïve retinal progenitor cells have two major developmental trajectories: one trajectory is toward ciliary margin cells, and the other trajectory is toward retinal neurons through a neurogenic state marked by *Atoh7* expression (see also Fig. 2A, 2K). In DKO retinas, the developmental trajectory toward ciliary margin cells is enhanced, whereas the developmental trajectory toward retinal neurons is defective. An ectopic trajectory lacking the Atoh7+ state leads to ectopic neurons.

## Discussion

In this study, we performed scRNA-seq of control and DKO eye cups at E13.5. We identified cell clusters based on their individual transcriptomes and then inferred developmental trajectories based on RNA velocities in the integrated control and DKO dataset. The UMAP graphing of individual cells revealed the relationships between tens of thousands of cells in eye cups. Differential expression analysis not only confirmed previous phenotype studies but also identified novel DEGs of cell clusters along developmental trajectories and DEGs caused by gene inactivation. We identified transcriptomes and cell trajectories that are jointly regulated by Six3 and Six6, providing deeper insight into the molecular mechanisms underlying the maintenance and differentiation of multipotent retinal progenitor cells.

Cell clustering and RNA velocity analysis reveal developmental trajectories of naïve retinal progenitor cells in the control retina. In the UMAP graph and streamline plot of RNA velocities, naïve retinal progenitor cells were progressed toward two major developmental trajectories: one trajectory was toward ciliary margin differentiation, and the other trajectory was toward neuronal differentiation. Cells at the end of the ciliary margin trajectory in the UMAP graph differentially expressed components of the Wnt signaling pathway, such as *Fzd1*, *Wls*, and *Wnt16*. Notably, *Fgf9* and *Fgf15* were expressed in naïve retinal progenitor cells in a pattern that was complementary to that of *Fzd1*, *Wls*, and *Wnt16* in the UMAP graph, forming opposing gradients in the Wnt and Fgf signaling. Additionally, ciliary margin cells did not express *Atoh7*. Relative positions of cell domains expressing *Fzd1*, *Wls*, *Wnt16*, *Fgf15*, and *Atoh7* in the UMAP graph faithfully reflect their relative positions on sections ^8, 37^. Additionally, ciliary margin cells were at an edge of naïve retinal progenitor cell cluster at G1 phase. These findings strongly indicate that ciliary margin cells are directly differentiated from naïve retinal progenitor cells at G1 phase under the regulation of the Wnt signaling, consistent with previous findings ^5, 7, 8^ .

Neuronal differentiation from naïve retinal progenitor cells goes through a neurogenic state marked by *Atoh7* expression. *Atoh7* expression overlapped the expression of naïve retinal progenitor cell markers at one end and early retinal ganglion cell markers at the other end in the UMAP graph, and thus *Atoh7* expression serves as a transition state toward neuronal differentiation from naïve retinal progenitor cells, consistent with recent findings by Mu lab ^20^. A few key gene markers for retinal ganglion cell differentiation, such as *Pou4f2*, *Isl1*, and *Sncg*, exhibited a progression of expression patterns toward the end of retinal ganglion cell trajectory in the UMAP graph. Key gene markers for photoreceptor, amacrine, and horizontal cell differentiation displayed expression patterns that were branched out from Atoh7+ neurogenic retinal progenitor cells in the UMAP graph. These findings indicate that neuronal differentiation from naïve retinal progenitor cells goes through a common neurogenic state marked by Atoh7 expression and then toward lineage-specific differentiation through a progression of cell states dictated by a cascade of regulators.

Cell clustering places Six3-deficient and Six3-wildtype DKO retinal cells into distinct regions in the UMAP graph, indicating drastic differences in their transcriptomes. Delineation of Six3- and Six6-deficient cells in the UMAP graph overcame the confounding issue that α-Cre is inactive in the central retina, enabling us to assess alterations of gene expression in retinal cells that were deficient for Six3 and Six6. Notably, an ectopic cell population was found in DKO retinas, strongly indicating the essential roles of Six3 and Six6 joint functions in the regulation of cell states and trajectories of the mouse retina.

Six3 and Six6 joint functions are required for the maintenance and proper differentiation of naive retinal progenitor cells. When Six3 was conditionally deleted in peripheral naïve retinal progenitor cells starting at E10.5 on top of Six6 germline inactivation, both naïve and neurogenic retinal progenitor cells exhibited deficits in marker expression: the expression of *Ccnd1*, *Sfrp2*, *Vsx2*, and *Atoh7* were significantly downregulated whereas *Neurod1* expression was significantly upregulated. One important aspect of Six3 and Six6 joint functions is to maintain naïve retinal progenitor cells in a proliferative state, as Six3 and Six6 dual deficiency caused downregulation of *Ccnd1*, fewer cells at S phase, more cells at G1 phase, upregulation of *Cdkn1c*, and DEGs that were enriched with GO terms related to cell cycles. Additionally, neuronal differentiation was disrupted whereas ciliary margin differentiation was enhanced. Notably, an ectopic cell population that displayed the upregulation of amacrine and forebrain markers was found. These findings indicate that naïve retinal progenitor cells have the developmental potentials for the neural retina and ciliary margins, and to some extent the forebrain. Six3 and Six6 are jointly required for the maintenance and proper differentiation of naïve retinal progenitor cells and the suppression of alternative cell fates such as the forebrain.

scRNA-seq enables us to identify DEGs caused by Six3 and Six6 dual deficiency in defined cell groups, a substantial advantage over bulk RNA-seq. We were able to identify DEGs caused by Six3 and Six6 dual deficiency in naïve retinal progenitor cells, neurogenic retinal progenitor cells, and the ectopic cell cluster, respectively. A significant number of DEGs identified by scRNA-seq, such as *Gadd45a*, *Sorb2*, *Rspo3*, *Enc1*, and *Dlx1* shown in feature plots, were not identified in previous bulk RNA-seq due to the lack of cell clustering ^7^. On the other hand, scRNA-seq missed out some DEGs identified by previous bulk RNA-seq due to the read depth. A union of DEGs identified by scRNA-seq and bulk RNA-seq revealed over 2000 candidate genes downstream of Six3 and Six6 joint functions. Six3 and Six6 jointly regulate this collection of target genes directly or indirectly during retinal differentiation.

Six3 and Six6 jointly balance the opposing gradients of Fgf and Wnt signaling in the maintenance and proper differentiation of naïve retinal progenitor cells. Previous studies indicate that opposing gradients of Fgf and Wnt signaling regulates the patterning of eye cups along the central-peripheral axis ^8^, but how these gradients are regulated is unclear. The downregulation of *Fgf9* and *Fgf15* and the upregulation of *Fzd1*, *Wls*, and *Wnt16* in Six3- and Six6-deficient cells establish that Six3 and Six6 jointly regulate opposing gradients of Fgf and Wnt signaling in the central-peripheral patterning of the mouse retina, enabling proper neuronal and ciliary differentiation.

Six3 and Six6 jointly repress the expression of actin-cytoskeleton regulators, including *Filip1* and *Ezr*. *Filip1*, which promotes the degradation of filamin A and therefore remodels the actin cytoskeleton ^35^, was one of the mostly up-regulated genes caused by Six3 and Six6 dual deficiency. The upregulation of *Filip1* and *Ezr* likely remodeled actin cytoskeleton, which explains the changes in cell shapes from elongated retinal progenitor cells to shortened ciliary margin cells upon Six3 and Six6 dual deficiency.

Target regulation by Six3 and Six6 joint functions is dependent on cell context. Some downstream genes, such as *Fzd1*, *Axin2*, and *Filip1*, displayed alterations of gene expression widely in Six3- and Six6-deficient cells. In contrast, *Wls*, *Rspo3*, *Wnt16*, and *Ezr*, exhibited changes in gene expression only in subsets of Six3 and Six6 deficient cells. These findings indicate that regulation of these downstream genes are dependent on cell context. The cell context could be related to their interacting proteins and chromatin states in individual loci. Further studies are needed to elucidate how Six3 and Six6 jointly regulate their target genes in a context-dependent manner.

Taken together, our scRNA-seq analysis of control and Six3- and Six6-deficient eye cups identifies cell states and trajectories in control retinas and discovers how these cell state and trajectories change upon Six3 and Six6 dual deficiency. Our scRNA-seq confirms previous phenotype analysis as well as provides novel insight into the mechanisms underlying early retinal differentiation. Six3 and Six6 jointly regulate the identities and developmental trajectories of multipotent retinal progenitor cells in the mouse retina.

## Acknowledgements

We are grateful to Dr. Roy Chuck for support, David Reynolds at the Genomics Core of AECOM for the preparation of single-cell libraries, and grants from NEI, NIH (R01EY022645 and R21EY029806 to W.L.) and Fight for Sight (FFS-PD-22-003 to R.S.).

## Materials and Methods

### Mice

*Six3* and *Six6* double knockout mice (*Six3^F/F^;Six6^−/−^;α-Cre*) were described in previous studies ^7^. The Animal Use Protocol was approved by the Institutional Animal Care and Use Committee at Albert Einstein College of Medicine.

### Cell capture from eye cups for scRNA-seq

Mouse embryos from the mating between *Six3^F/F^;Six6^+/−^;α-Cre* male and *Six3^F/F^;Six6^+/−^* female were harvested at E13.5. Intact eye cups containing the neural retina and lens were dissected out from neighboring tissues. Eye cups from an embryo with the genotype of *Six3^F/F^;Six6^−/−^;α- Cre* were identified based on GFP+ rosettes in the retinas under a stereo fluorescence microscope. The correlation between GFP+ rosettes and the genotype of *Six3^F/F^;Six6^−/−^;α-Cre* was previously established in pilot studies. Eye cups from an embryo without GFP were used as a control. Tails of the selected embryos were collected for genotyping. Then, eye cups were dissociated into single cells using activated Papain (Worthington Biochemical) for cell capture using the 10x Chromium fluid device, targeting 10,000 cells for each sample. Genotyping of the embryos confirmed the genotype *of Six3^F/F^;Six6^−/−^;α-Cre* for DKO retinas and the genotype of *Six3^+/−^;Six6^+/−^* for the littermate control. Previous phenotype analysis indicates that embryos with the genotype of *Six3^+/−^;Six6^+/−^* are indistinguishable from wildtype embryos ^7^. Captured cells were used for library preparation using the 10x Genomics Single Cell 3’ kit (version 3).

### Analysis of scRNA-seq

The libraries of DKO and control retinas were sequenced on a Hiseq lane (GeneWZ). Sequencing reads were mapped to mm10 (3.0.0) using CellRanger (3.1.0).

Quality control and data analysis were performed using the Seurat package (3.2.0) ^26^. Briefly, cells were filtered (nFeature_RNA > 200 & nFeature_RNA < 6000 & percent.mt < 20), resulting 9822 and 12146 cells for control and DKO cells, respectively. The two datasets were normalized, and their variable features were found. Anchor cells of the two datasets were identified for integration using the function *FindIntegrationAnchors*, and the two datasets were integrated using the function *IntegrateData*. The integrated dataset was scaled using the function *ScaleData*. Dimension reduction, graph-based visualization, and clustering were accomplished using functions *RunPCA*, *RunUMAP*, *FindNeighbors*, and *FindClusters*. DEGs were identified using the function *FindMarkers*. Functional annotations of DEGs were achieved using the function *enrichGO* in the clusterProfiler package ^38^. The cell trajectory analysis of retinal cells was done using the scVelo package ^28^.

### In situ hybridization

Mouse embryos from the mating between *Six3^F/F^;Six6^+/−^;α-Cre* male and *Six3^F/F^;Six6^+/−^* female were harvested at E13.5, fixed in 4% paraformaldehyde in cold overnight, and then soaked in 30% sucrose for cryopreservation. DKO and control embryos were sectioned at 10 μm for in situ hybridization using DIG-labeled RNA probes (Sigma). DNA templates for making RNA probes were generated using PCR with primers (Suppl. Tab. S3).

### Statistical analysis

Default settings of p value and percent of cell population in the function *FindMarkers* were used the identification of DEGs. For GO term enrichment analysis, DEGs were further filtered with padj. < 0.01. Results of in situ hybridization represented a minimum of three embryos.

## Data availability

Fastq files of scRNA-seq were deposited in the GEO database (GSExxx).

## Legends for supplemental figures

**Supp. Fig. S1. Identification of non-retinal cell clusters in the integrated scRNA-seq dataset.** Related to Fig. 1. **(A-D)** Clusters 8 and 19 differentially express *Hba-x* (A, B); clusters 8, 11, and 19 differentially express *Hba-a1* (A-D). These findings indicate that clusters 8, 11, and 19 are red blood cells. **(E-H)** Cluster 17 differentially express *C1qb* and *Tyrobp*, indicating that these cells are white blood cells. **(I-L)** Clusters 12 and 14 differentially express *Cryab* and *Cryaa*, indicating that they are lens cells. R, red blood cells; W; white blood cells; L, Lens cells.

**Supp. Fig. S2. Clusters 2 and 13 are marked by negative gene markers that are also nearly absent in red blood cells and differentiating lens cells.** Related to Fig. 1. **(A-H)** Clusters 2 and 13 barely express *Kcnq1ot1*, *mt-Co1*, *mt-Nd2*, and *mt-Nd4*. These gene markers are also nearly absent in clusters 8, 11, 19, and 14.

**Supp. Fig. S3. Violin plots of nCount_RNA, nFeature_RNA, and percent.mt for cell clusters.** Related to Fig. 1. **(A)** Clusters 2, 13, and 14 have lower values for nCount_RNA. **(B)** Clusters 2, 8, 11, 13, and 14 have lower values for nFeature_RNA. **(C)** Clusters 2, 8, 11, 13, and 14 have lower values for percent.mt.

## Legends for supplemental tables

**Supp. Tab. S1. Proportions of cell clusters and assigned cell types.** Related to Fig. 1.

**Supp. Tab. S2. Proportions of cells at three cell cycle phases.** Related to Fig. 1. Non-retinal cells were removed before the counting.

**Supp. Tab. S3. Sequences of PCR primers for the generation of DNA templates for making RNA probes.** Related to Fig. 7.

