## Supplementary figures and images for "Six3 and Six6 jointly regulate the identities and developmental trajectories of multipotent retinal progenitor cells in the mouse retina"

### supplemental figures

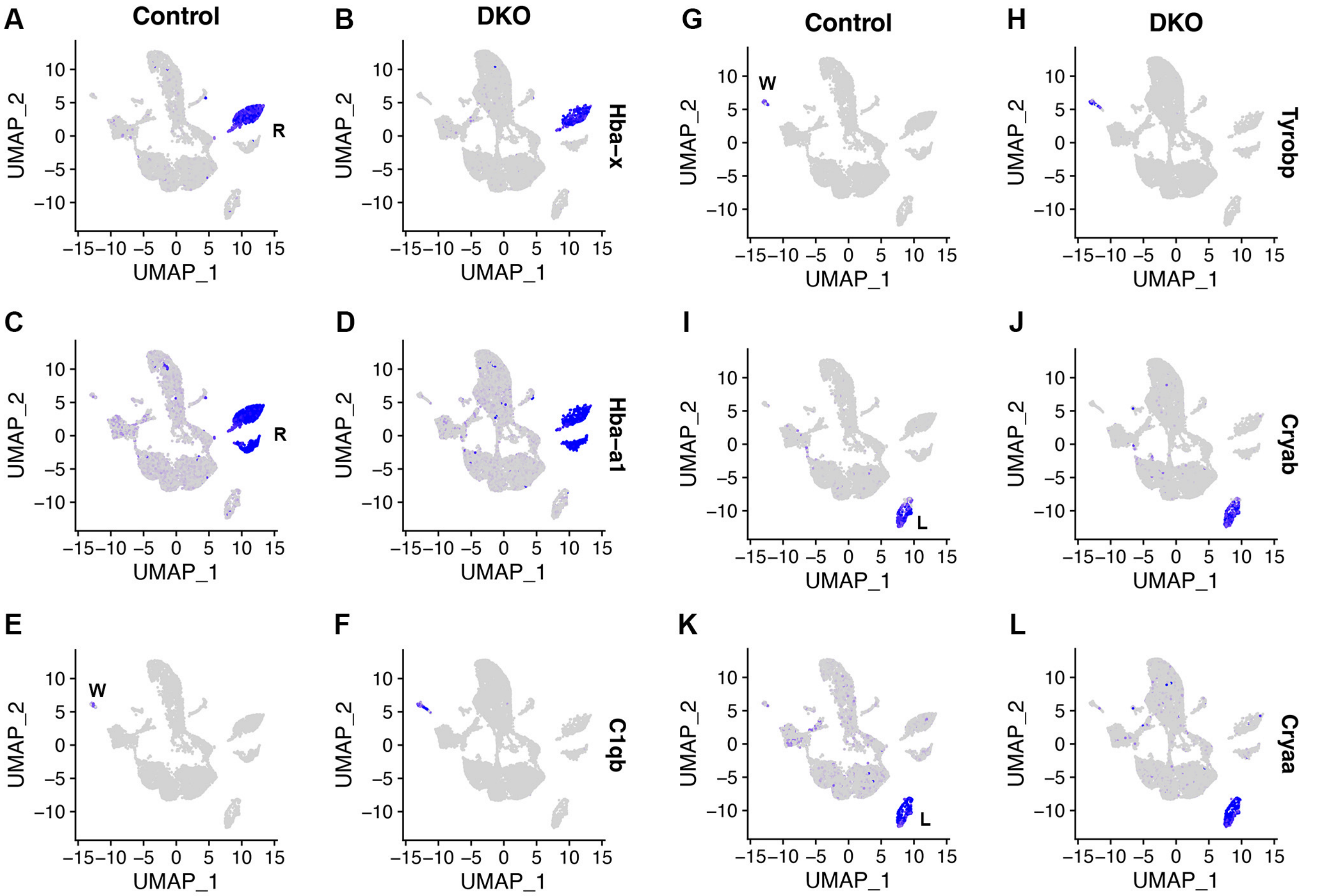

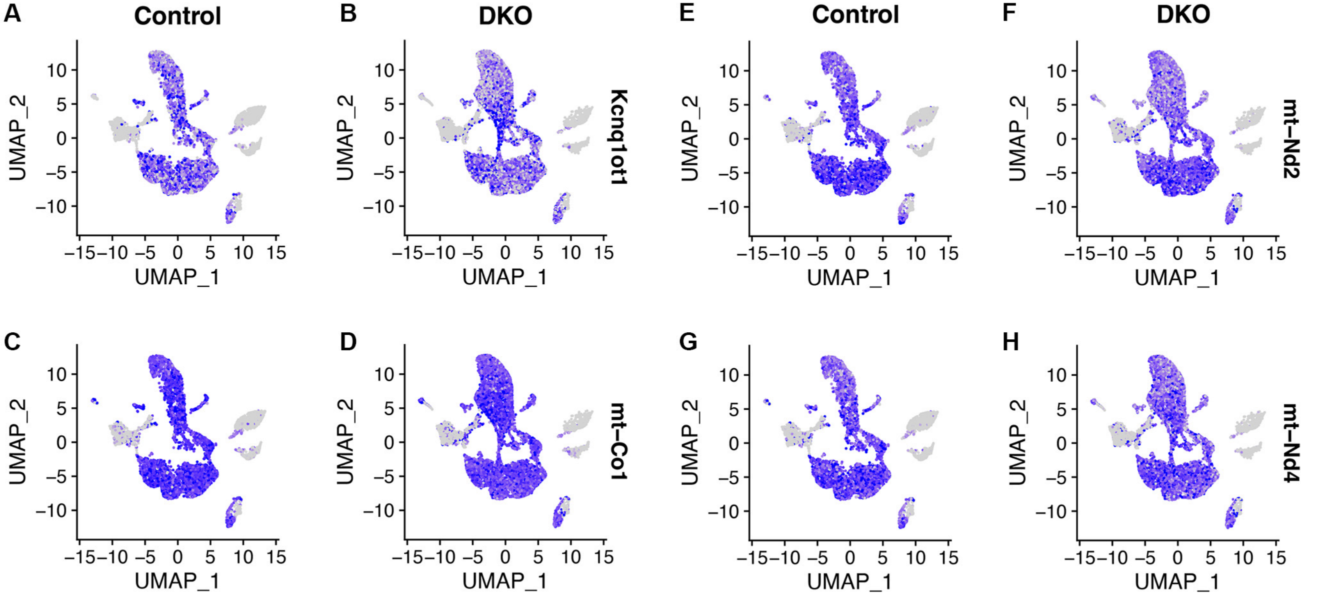

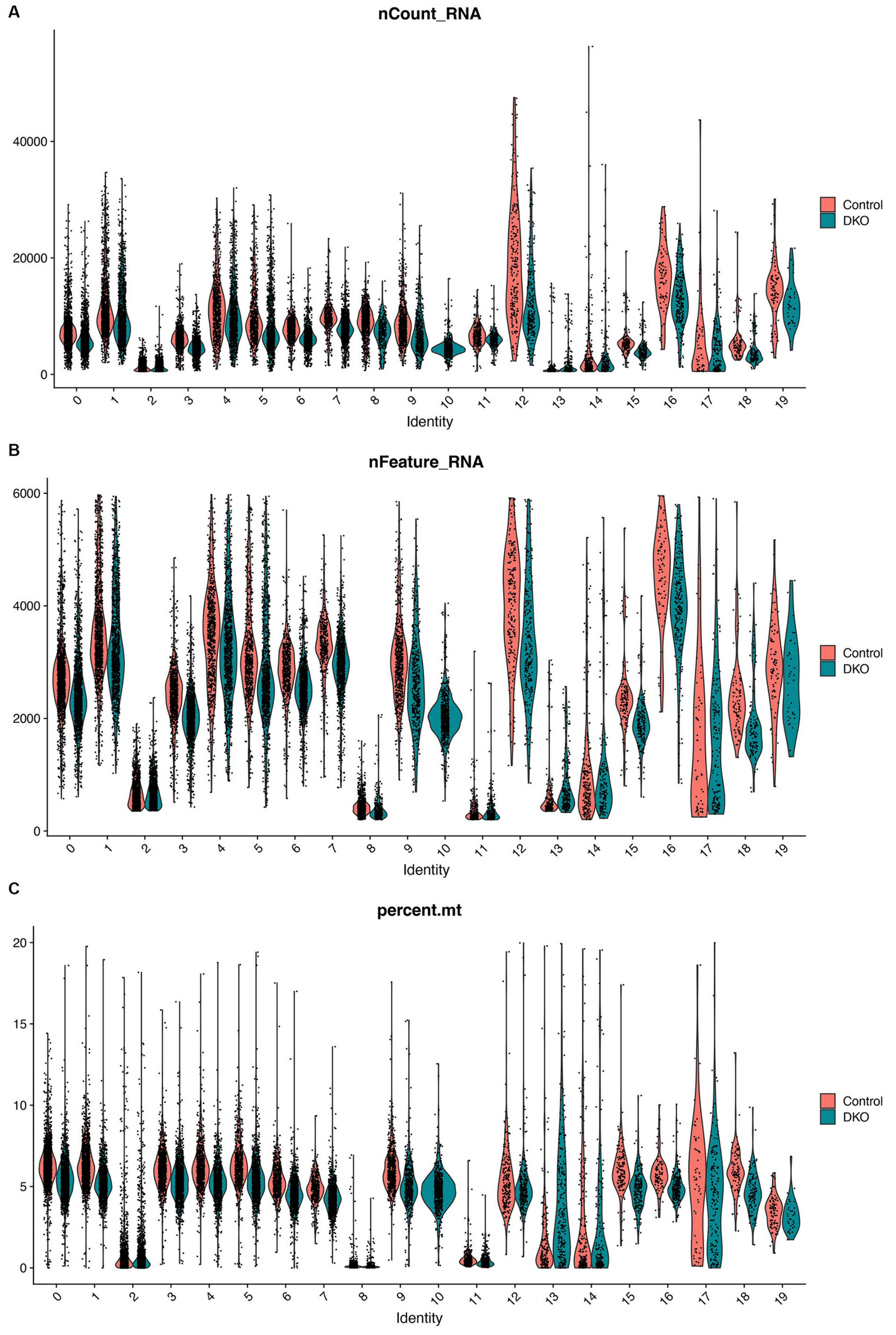
