## Supplementary material for "Six3 and Six6 jointly regulate the identities and developmental trajectories of multipotent retinal progenitor cells in the mouse retina": supp. Tab S1

| Cluster | Ctrl | DKO | Assigned Cell Types |
|---|---|---|---|
| 0 | 0.161 | 0.118 | Naïve RPC |
| 1 | 0.133 | 0.096 | Naïve RPC |
| 2 | 0.107 | 0.095 | RPC with fewer genes; Six3+ regions |
| 3 | 0.083 | 0.103 | Advanced neurogenic RPC + Early RGC + Ectopic cell |
| 4 | 0.073 | 0.080 | Naïve RPC |
| 5 | 0.068 | 0.084 | Naïve RPC_G1 including CM |
| 6 | 0.057 | 0.088 | Early RGC + Advanced neurogenic RPC + Ectopic cell |
| 7 | 0.037 | 0.087 | Advanced RGC, Six3+ regions |
| 8 | 0.083 | 0.024 | Red blood cell |
| 9 | 0.064 | 0.037 | Early neurogenic RPC |
| 10 | 0.000 | 0.051 | DKO-specific |
| 11 | 0.027 | 0.022 | Red blood cell |
| 12 | 0.022 | 0.021 | Lens cell |
| 13 | 0.018 | 0.020 | RGC with fewer genes, Six3+ regions |
| 14 | 0.022 | 0.013 | Lens cell with fewer genes |
| 15 | 0.013 | 0.016 | Amacrine + Horizontal |
| 16 | 0.008 | 0.020 | RGC_16 |
| 17 | 0.006 | 0.013 | White blood cell |
| 18 | 0.009 | 0.010 | Photoreceptor / Cone |
| 19 | 0.009 | 0.003 | Immature red blood cell |
