## Supplementary material for "Six3 and Six6 jointly regulate the identities and developmental trajectories of multipotent retinal progenitor cells in the mouse retina": Supp. Tab. S3

| ID | Sequence | Gene | Size |
|---|---|---|---|
| RD43 | cctgcacagccagcattttt | Gja1 UTR #F Mouse In Situ Probe | |
| RD44 | **GAG**taatacgactcactatagggcatttaccagcaccgggact | Gja1 UTR #R Mouse, **RNA Polymerase Binding Site**/T7 Polymerase | 814bp |
| RD47 | GTCCGCTTCTTCTGGTGAGT | Dct CDS #F Mouse In Situ Probe | |
| RD48 | **GAG**taatacgactcactatagggaccgtggtgaatgacccaa | Dct UTR #R Mouse, **RNA Polymerase Binding Site**/T7 Polymerase | 824bp |
| RD51 | agcctcttcccctggagtag | Wls UTR #F Mouse In Situ Probe | |
| RD52 | **GAG**taatacgactcactatagggcagcacctgggtactccaag | Wls UTR #R Mouse, **RNA Polymerase Binding Site**/T7 Polymerase | 845bp |
| RD177 | CCTGCTCAGAACGCCAGAA | Rspo3 mRNA #F Mouse, In Situ Probe | |
| RD178 | **GAG**taatacgactcactatagggTTCTGGGCAACTGTCAAGGC | Rspo3 mRNA #R Mouse, **RNA Polymerase Binding Site**/T7 Polymerase | 894bp |
